# Environmental DNA monitoring of waterfowl reveals community changes during migration

**DOI:** 10.1101/2025.11.11.687912

**Authors:** Luciana Guimaraes de Andrade, Steve Michael Bogdanowicz, Holger Klinck, David Lodge, Jose A. Andrés

**Author notes:** These authors contributed equally to this work (Luciana Guimaraes de Andrade, Jose A. Andrés). These authors contributed equally to this work (Steve Michael Bogdanowicz, Holger Klinck, David Lodge).

## Abstract

Large-scale visual surveys are an integral part of waterfowl conservation and management programs. In this paper we explore the possibility of using environmental DNA-based surveys as a cost-efficient, complementary tool to estimate populations of North American waterfowl species. To achieve this, we first evaluated the performance of all currently available avian metabarcoding primers and compared them to newly designed primers targeting the mitochondrial ND2 gene within the Anatidae tribes of North America. All the existing avian assays showed strong cross-priming amplification with other vertebrates. In contrast, in-silico analyses of our waterfowl targeted assays showed a high degree (>90%) of avian specificity, encompassing all the 132 Anatidae species sequenced thus far. We used this targeted metabarcoding approach to track the temporal variation in the relative abundance of waterfowl species during the fall migration at Montezuma National Wildlife Refuge, New York, a major resting area for waterfowl on their journey to and from North American nesting areas. We compared eDNA results with visual surveys conducted by us and from those reported on eBird. Our results showed that eDNA detected all waterfowl species (n= 25) during fall migration. Positive correlations existed between standardized eDNA read counts and the relative abundance of waterfowl species as reported in eBird on the day of sampling and up to five days prior. However, this approach did not provide a good metric for absolute abundance of waterfowl species: only 8 out of 25 waterfowl species showed significant correlations between the number of eDNA reads and the total abundance of birds. Overall, while eDNA-targeted metabarcoding has not yet been applied to study bird communities extensively, our results demonstrate that this technique can be used as an effective complementary tool for assessing species composition of waterfowl communities and estimating relative abundance of species within those communities.

## Introduction

Large-scale wildlife monitoring programs play a crucial role in addressing the management and conservation objectives for waterfowl, a taxonomic group that is ecologically, economically, and culturally significant [45]. To estimate population sizes and trends, traditional waterfowl monitoring efforts cover extensive geographic areas using a combination of visual surveillance techniques [5, 6, 16, 21, 35, 51, 67, 74]. Visual surveys conducted from the ground are labor-intensive and constrained by limited access to private lands [17]. Thus, aerial surveys have become the preferred tool in these large-scale surveys (e.g., [76]). Despite their advantages over ground counting methods, aerial surveys also have important limitations: depending on the sampling design, generalization to unsampled areas may be difficult. In addition, aerial surveys may overlook regions of high waterfowl use if distributions change over time [35]. Additionally, the infrequent nature of many aerial surveys does not allow for accurate estimation of abundance variation among survey periods, and analyzing aerial images is time-consuming even with the use of machine learning techniques [21, 28, 38, 58].

Considering the above limitations and the disturbance caused by low-altitude flying [26], alternative, cost-effective tools for monitoring waterfowl populations are needed. High-throughput sequencing of macro-organismal DNA from environmental samples, known as eDNA metabarcoding, has proven to be a robust and efficient technique to characterize biological communities across a diverse array of aquatic taxa (e.g., [2, 7, 18, 27, 32, 52]. Several lines of evidence suggest that extending eDNA monitoring of waterfowl populations should be feasible. First, eDNA metabarcoding surveys using fish and mammal primers typically “bycatch” bird species [34, 48, 49, 57, 60, 73, 81]. Second, analyses of water samples from zoo bird cages and natural environment suggest that universal avian metabarcoding primers can be used to describe bird communities [19, 75]. Third, eDNA assays have detected the true geese and swan species in both zoo ponds and natural lakes [37]. While this technique has not yet been systematically applied to survey bird communities [41], eDNA detections compared favorably with visual bird surveys in a recent study in a Chinese lake [64]. However, other bird eDNA studies have faced challenges related to primer selection, sampling strategies and estimating the abundance of bird communities [40, 56, 64].

In this study we aim to take the first step towards the future implementation of robust eDNA-based waterfowl biosurveillance programs by answering two key questions: Can a full, diverse waterfowl community be reliably detected from water samples; and are waterfowl abundance patterns inferred from water samples consistent with those obtained from intensive ground visual surveys?

To answer these questions we first tested a series of published bird metabarcoding primers [4, 22, 42, 43, 55, 65, 66, 74]. Although most of these universal assays successfully amplified waterfowl eDNA, our initial results showed PCR primer bias and cross-amplification with other abundant taxa, including fish, severely affected the sensitivity of the assay, increased false-negative rates, and biased abundance estimates. To minimize these limitations we designed a metabarcoding assay [*sensu* 77] specifically designed to detect North American waterfowl species. We then examined the applicability of this method by tracking the temporal variation in relative abundance of waterfowl species during fall migration at Montezuma National Wildlife Refuge, New York, a major resting area for waterfowl and other waterbirds on their journeys to and from North American nesting areas.

## Materials and Methods

### Study areas

Our primary study area was the Main Pool at Montezuma National Wildlife Refuge, in Seneca County, New York, United States (Fig 1).

**Fig 1.**
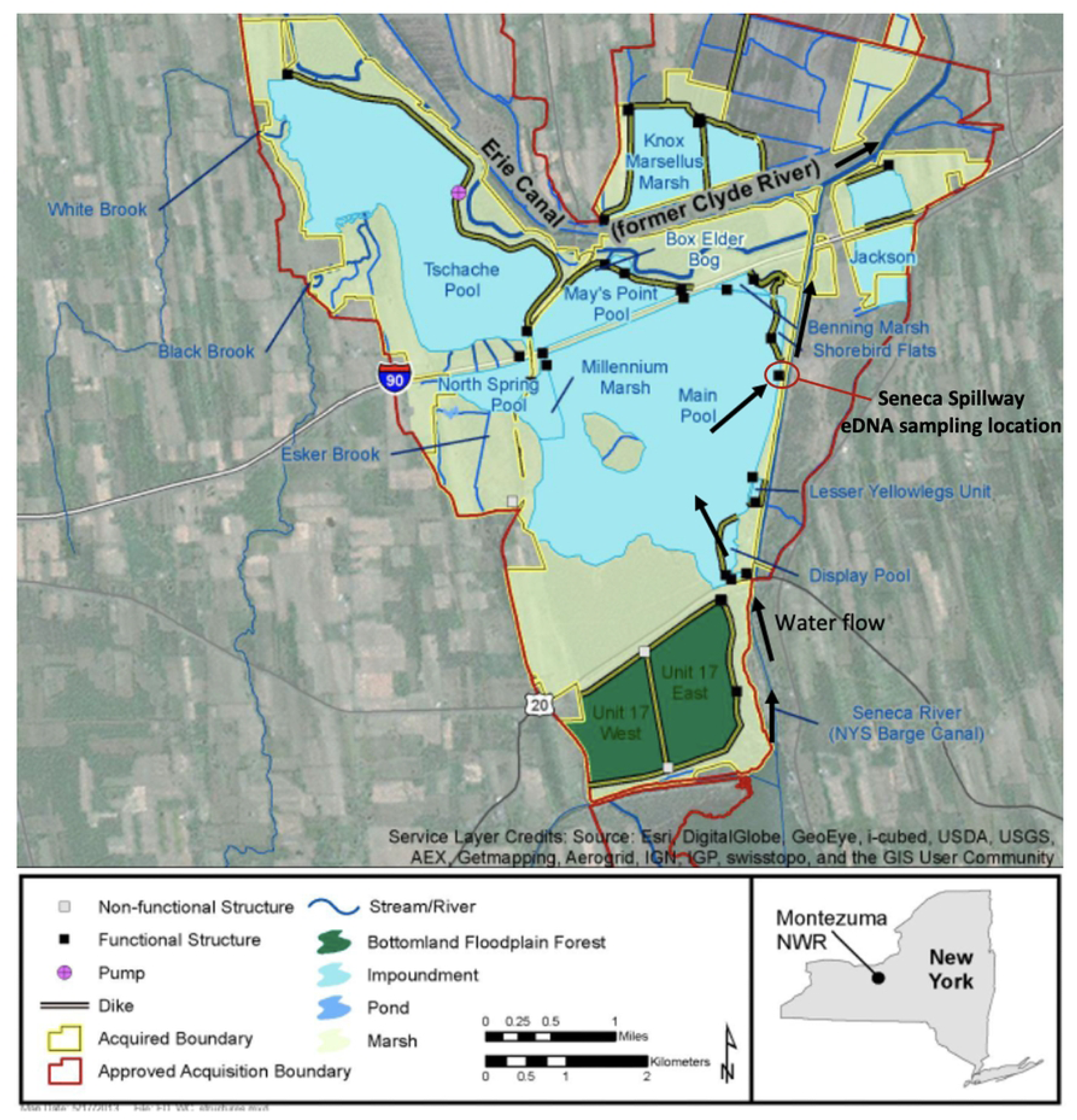
Water flow in the Main Pool at Montezuma NWR modified from [78] to show the Seneca Spillway where water samples were taken in 10 consecutive weeks during the peak of the 2020 waterfowl fall migration. Black arrows represent the general direction of the water flow during this period.

Montezuma NWR is a 4,000-hectare wetland complex located on a major migratory corridor in the Atlantic Flyway, sustaining up to a million waterbirds during non-breeding seasons [10]. The Main Pool (also referred to as Black Lake on some maps) is a shallow 309-hectare impoundment, averaging about 60 cm deep, whose water levels are managed to provide emergent marsh and other habitat types for migratory birds and resident wildlife. The water level was 1 m during our field sampling, from September to November 2020. Other than precipitation, the primary intake for the Main Pool is water from Cayuga Lake, which is provided through a ditch with a gravity-fed connector [78]. Water leaves the Main Pool via the Seneca Spillway outlet, which empties into the Cayuga-Seneca Canal. The overall flow of water is from Cayuga Lake northward through the Main Pool to the Cayuga-Seneca Canal (Fig 1). To conduct the initial assessment of both general and targeted bird metabarcoding primers, we collected water samples from both Sapsucker Woods Pond in Tompkins County, New York and two water bodies at the Huyck Preserve (Lincoln Pond and Lake Myosotis) in Albany County, New York (S1 and S2 Figs). Then, to test the validity of targeted metabarcoding as a monitoring tool, we took weekly water samples (n=3 + 1 field blank) at Montezuma NWR during the peak of the 2020 fall waterfowl migration (September 24 -November 29). All weekly water samples were taken near the Seneca Spillway outlet (Fig 1).

### Water sampling

At each sampling location, we sampled eDNA by collecting and filtering 250 ml water samples and one distilled water negative control. Water was collected by submerging a sterilized wide-mouth Nalgene container (Thermo Fisher Scientific) < 0.1 m beneath the water’s surface following the recommendations of [31] to minimize the probability of contamination. Each of the samples was then filtered through a 25 mm diameter cellulose-nitrate filter, 1.0 µm pore size, (Whatman, GE Healthcare) using a luer-lok sterile syringe system. Filtration blanks of distilled water were processed in the same way. Filters from each sample were immersed in 700 μl Longmire’s buffer, transferred to the lab within 24 h, and stored at −80°C until DNA extraction. Water samples described above were taken just after our visual surveys.

### Visual surveys

From the shore, we conducted exhaustive visual surveys of all waterfowl on the Montezuma NWR Main Pool during weekly visits from September 24 to November 29, 2020. These ground-based surveys were performed along a 1.3 km stretch of the Montezuma NWR Wildlife Drive, immediately adjacent to the Main Pool, using Zeiss 10 x 40 binoculars and a Swarovski spotting scope with a 20–60x eyepiece. The average survey start time was 9:28 am (range = 7:47 am to 10:33 am) and the average survey duration was 131 minutes (range = 92 to 149 minutes). To supplement our ground visual surveys, we compiled waterfowl observational data from eBird, the world’s largest curated database of citizen science bird sightings [72], for each water sampling date plus a five-day look-back period. We followed best practices for using eBird data [71], utilizing only complete checklists (i.e., all the birds observed are reported) that followed eBird’s “Stationary” or “Traveling” protocols. For dates with more than one complete eBird checklist, we calculated species abundance as the mean number of individuals from all complete checklists.

### Avian metabarcoding primer tests using natural eDNA

Based on previous bird eDNA and phylogenetics studies we tested 11 unique primer pairs targeting different fragments of 5 mitochondrial loci: cytochrome c oxidase I [42, 55], 12 S rRNA [22, 74], NADH dehydrogenase subunit 2 [65, 66], and Cytochrome b [4,43] (S1 Table). Insert length and amplicon size for each genetic marker were presented in S2 Table. The performance of each primer pair was tested on the eDNA samples from Sapsucker Woods Pond and Huyck Preserve (see above).

### Anatidae primer design

Based on the results of our avian metabarcoding tests, and the availability of reference sequence data we chose the mitochondrial gene ND2 as a molecular marker for waterfowl species. We thus downloaded all ND2 gene sequences from NCBI corresponding to all North American Anatidae tribes as of May 27, 2020. Subsequently, we selected up to two DNA sequences for each species from distant geographic locations. The 85 selected sequences representing 54 species and 19 genera in the family Anatidae were aligned using the ClustalW algorithm. The manual inspection of the alignment did not reveal suitable ND2 regions for a single “universal” set of primers for Anatidae. Thus, we followed a strategy similar to that of [15] to generate a Tamura-Nei NJ tree to identify groups of similar sequences sharing relatively conserved regions (S4 Fig). For each of the three groups we found, we designed a single set of PCR primers using the software Geneious Prime 2020.1.1 (Table 1).

**Table 1.**
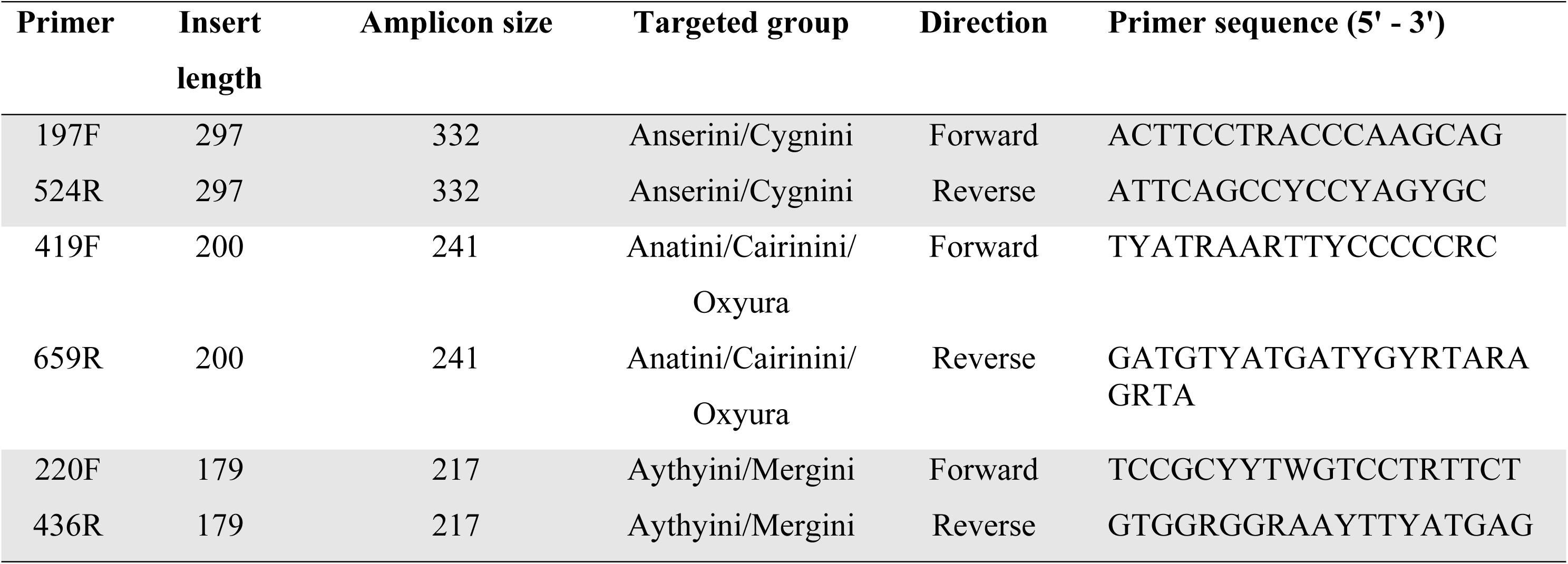
ND2 primers developed for DNA amplification of waterfowl species. Insert length, amplicon size and primer sequence for each waterfowl primer were included.

### In-silico assessment of Anatidae ND2 primers

The performance of the primers was tested as follows. First, we used CRABS [39] to generate a custom reference database including all ND2 vertebrate sequences found in the NCBI nr repository (n=157,730). Using the same pipeline, we performed in-silico PCR to produce a list of unique metabarcodes and assessed the specificity of the primers by calculating the percentage of Anatidae species amplified over non-target organisms using a maximum mismatch of 2 nucleotides in each primer.

### DNA extraction and library construction

All DNA extractions were carried out in a dedicated eDNA laboratory at Cornell University (Ithaca, NY), with UV light treatment overnight, HEPA filtered air under positive pressure, personnel full body suits, breathing masks and face cover shields. We used the DNeasy Blood and Tissue extraction kits (Qiagen Inc.) and the protocol described in [68]. All lab reusable materials, such as forceps, containers, and the lab counters, were regularly sterilized with 50% bleach solution and treated under UV light before and after each round of DNA extractions. Extraction negative controls were included in each of the eDNA extractions rounds. Extracted samples were placed in a −20^°^C freezer until library preparation. First-stage eDNA amplification was conducted in triplicate for each sample and negative control (i.e., field, filtration, extraction, and PCR blanks) in PCR volumes of 20 μl containing 8.475 μl molecular biology grade (MBG) water, 5 μl of 5x OneTaq Standard Reaction buffer, 0.2 μl of 10 μM primer (forward and reverse), 0.5 μl of 10mM dNTPs, 0.125 μl of One Taq Hot Start polymerase, 2.5 μl of bovine serum albumin (0.4 μg/l), and 3 μl of eDNA extract as template. PCR cycling for each waterfowl ND2 primer pair was described in the supplementary material.

PCR products were then pooled by sample and tagged in a second-stage PCR using Illumina Nextera N7 and N5 indexes. The second PCR was carried out with a 20-mL reaction volume per eDNA sample containing: 3 μl DNA, diluted with MGB water 1:1, 10.9 μl MGB water, 4 μl 5x OneTaq buffer, 0.4 μl 10mM dNTPs, 0.8 μl each of 10 mM Nextera N7 and Nextera N5, and 0.1 μl OneTaq Hot Start DNA polymerase (New England – BioLabs Inc.).

PCR was run using the following thermal cycling parameters: six cycles of 30 seconds denaturation at 95°C, 1 minute annealing at 62°C, and 1 minute extension at 68°C, with a final extension for 10 minutes at 68°C. The resulting amplicons were purified and size-selected using Agencourt AMPure XP beads (reaction ratio AMPure beads 0.9: PCR product 1; Beckman Coulter Genomics), and the concentration of each library was estimated using the Qubit dsDNA Broad Range Kit and Qubit 2.0 fluorometer (Life Technologies). We pooled the barcoded PCR products for Anatini and Aythyini primers and all the libraries were diluted to 2 nM for paired-end sequenced V2 (2 x250bp) on a Miseq platform (Illumina, Inc.).

### Bioinformatic analyses

Initial quality control of raw sequence data was performed using FastQC/MultiQC [3]. Degenerate bases and adaptors were trimmed from demultiplexed paired reads and sequences 35 bp in length were removed using Trimmomatic 0.33 [11]. Processed sequences were then analyzed using DADA2 1.16 [13], which involved removing forward and reverse primers, trimming reads to 200 bp for Anatini/Aythyini primers, and 390 bp for Anserini/Cygnini, and discarding sequences with expected number of errors >2 (EE, calculated based on Phred scores). The sequences were then denoised using the DADA2 error filtering algorithm. Forward and reverse reads for Anserini/Cygnini primers were merged with a minimum overlap of 20 bp and a maximum of one accepted error in the overlap region, and putative chimeras were removed. For Anatini/Aythyini primers, because of the small size of the amplicon (200 bp and 179 bp respectively), only forward reads were used for subsequent analysis. The resulting dataset contains amplicon sequence variants (ASVs) and the associated read counts for each sample. Taxonomic assignments for the resulting amplicon sequence variants (ASVs) were obtained using the BLASTn algorithm [14] and the nucleotide and taxonomy (nt/taxdb) databases from NCBI [9, 23]. We retained the top five target sequence matches for each ASV and assigned species-level taxonomy to ASVs that matched a single species with a sequence identity of ≥98%. If multiple species matched the query sequence equally well, ASVs were assigned the lowest common taxonomic rank (genus or family) among the target sequences with equal percent identity. ASVs that could not be classified to the family level were excluded from further analysis. ASVs with taxonomic assignments were filtered by removing ASVs with read counts fewer than the average number of non-zero reads summed across the DNA extraction and PCR negative controls.

### Statistics

While abundant waterfowl species can be found in the tens of thousands of individuals, rare species might be represented by only a handful of individuals. To account for abundances that differed across species by five orders of magnitude for both visual surveys and ASVs, we used log-log correlations to reflect the relationship between number of ASVs and the visual estimates of bird number, as previous quantitative eDNA metabarcoding studies have done [50, 69]. Because waterfowl daily density is likely to exhibit short-term temporal autocorrelation [1] and because the rate of eDNA degradation can vary from hours to weeks depending on the ecosystem, target species, and eDNA capture method [47], we calculated the correlations between the number of ASVs and individuals, as well as between the number of birds/species (from eBird lists) on different dates, up to 5 days prior to each eDNA sampling event. Then, to visualize the temporal covariation of waterfowl and their associated eDNA reads during migration, we constructed a correlation-based network (CNA) including eDNA and visual count datasets. In the CNA, the nodes represent dates during the fall migration and the edges between them the significant, positive, log-log correlation coefficients (≥ 0.6) between visual surveys or between visual surveys and the standardized number of DNA reads obtain by combining the Anserini/Cygnini and Anatini/Aythyini datasets.

To compare waterbird relative abundance estimates based on visual surveys with those based on eDNA read counts we focused on sampling dates with at least three independent complete eBird checklists (September 24th, October 15th, November 7th, and November 29th). For each of these dates, we compared the log-log correlation coefficient between our visual observations and eDNA counts to the log-log correlation coefficient between our visual observations and eBird data. All statistical analyses were performed with R.4.0.4 [59] and igraph.

## Results

### Evaluation of avian metabarcoding primers on eDNA samples

The pilot experiments at Sapsucker Woods Pond and the two Huyck Preserve waterbodies revealed a high level of cross-amplification of non-targeted taxa for all previously published avian metabarcoding primers. COI primers showed very little specificity, amplifying mostly bacteria (72% of reads) and fishes (20% of reads) (S3 Fig). Similarly, CytB primers amplified mostly other taxonomic groups (fish 54% of reads; mammals 44% (S3 Fig). 12S and ND2 primers amplified mostly avian species (86% and 78% of reads respectively), but the majority of reads were represented by only one waterfowl species (Canada goose) that was the predominant species in the two locations during the field sampling (S3 Fig). 12S and ND2 also cross-amplified teleosts (12S: 8% of reads; ND2: 20% of reads), potentially compromising the ability to detect waterfowl species in water bodies where fish species are abundant.

### In-silico and experimental evaluation of waterfowl-targeted primers

To avoid fish cross-amplification and increase waterfowl detection, we designed three pairs of primers, each of which targeted groups that corresponded to Anatidae tribes (Table 1).

In silico-PCR simulated against all vertebrate sequences available on the nt NCBI repository revealed that the targeted primers could amplify all ND2 waterfowl sequences available, encompassing 76% (132/174) of the species recognized by the IOC World Bird List International Ornithological Committee [29]. None of the targeted primers are waterfowl-specific when considering a maximum of two mismatches between each primer and the template. However, within these parameters the newly designed primers exclusively amplify bird species (S5 Fig). The experimental evaluation in two water bodies with contrasting waterfowl species diversity indicated high sensitivity and accuracy. At the Huyck Preserve, where visual surveys indicated low abundance and diversity of waterfowl (Canada goose, n=406; Wood duck, n=56; hooded merganser, n=5; mallard, n=11; American black duck, n=1), we generated approximately 4 million high-quality reads, identifying 46 ASVs corresponding to four of the five observed species, with the Hooded Merganser being the only one not detected. Additionally, we identified DNA from three species that were not visually observed: American Wigeon, Northern Pintail, and Redhead. At Montezuma NWR, where 25 waterfowl species were reported during our sampling period (Table 2), the eDNA dataset comprised over 11 M high-quality sequence reads (∼650,000 reads/sample) corresponding to 322 waterfowl ASVs. eDNA successfully detected all waterfowl species present in Montezuma NWR, whose abundance spanned 5 orders of magnitude (from less than 10 to over 5,000 individuals). While most of the ASVs could be undoubtedly assigned to a single species, the targeted ND2 fragments could not be distinguished three species pairs: mallard/American black duck, tundra swan/trumpeter swan, and greater/lesser scaup. Accordingly, both visual and DNA read counts for these species pairs were pooled for subsequent analyses.

**Table 2.**
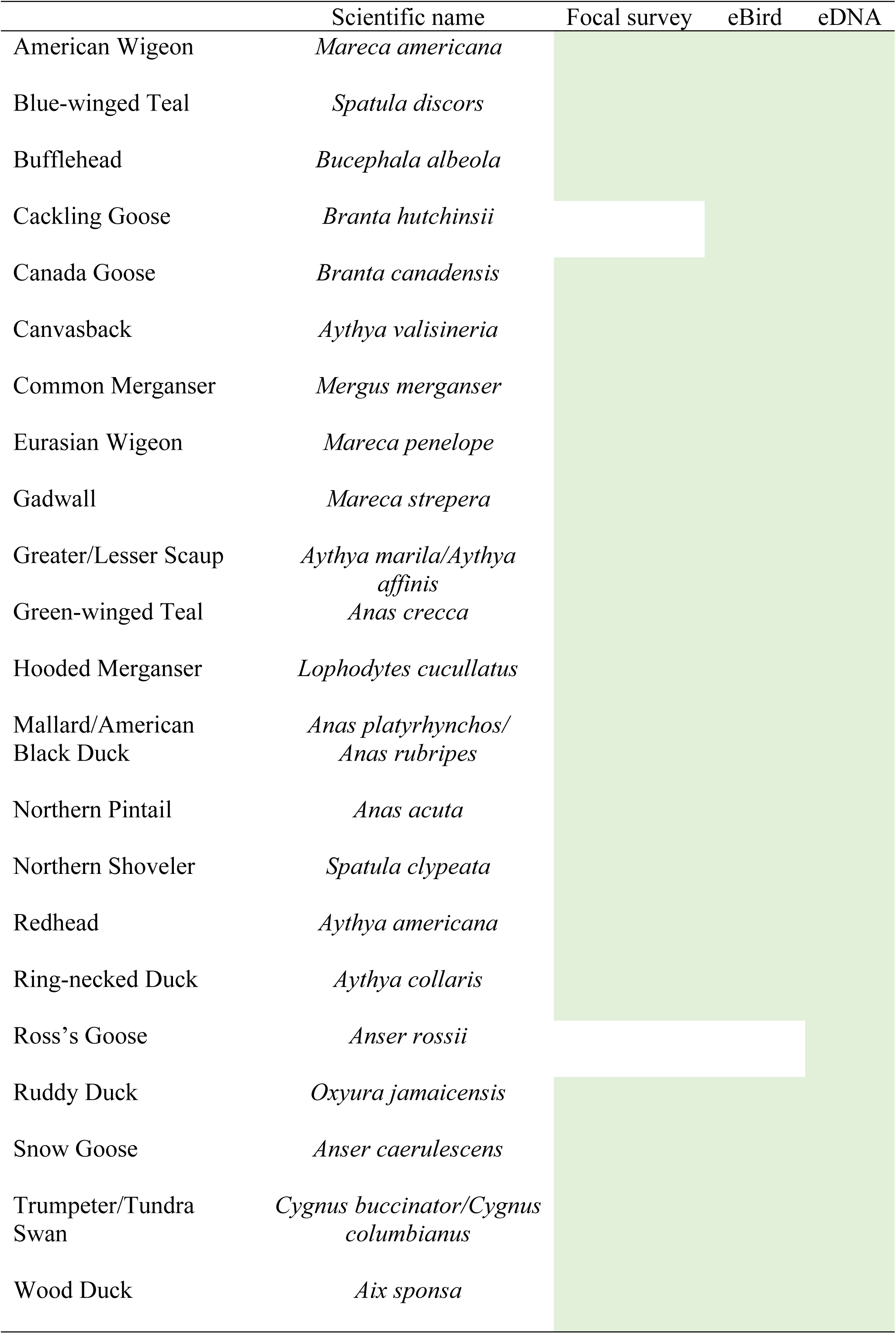
Waterfowl species detected in the Main Pool, at Montezuma NWR, by our ground visual surveys, eBird surveys, and eDNA during field sampling in 2020.

### Temporal variation in the abundance of waterfowl species

Overall, there was a weak correlation between the number of eDNA reads and the total abundance of birds, as determined by the number of individuals observed during the visual survey on the day of sampling (Fig 2). Only 8 out of 25 species (including blue-winged teal, bufflehead, canvasback, green-winged teal, and wood duck) showed a statistically significant correlation (P_one-tail_> 0.05). For example, Canada goose eDNA reads were consistently overrepresented, while ring-necked duck reads were underrepresented. Although both species exhibited strong seasonal variation in the number of individuals, there was little seasonal variation in their eDNA read counts. Thus, the observed variation in the standardized number of eDNA reads did not accurately reflect the temporal fluctuations in the absolute abundance of waterfowl species observed during fall migration.

**Fig 2.**
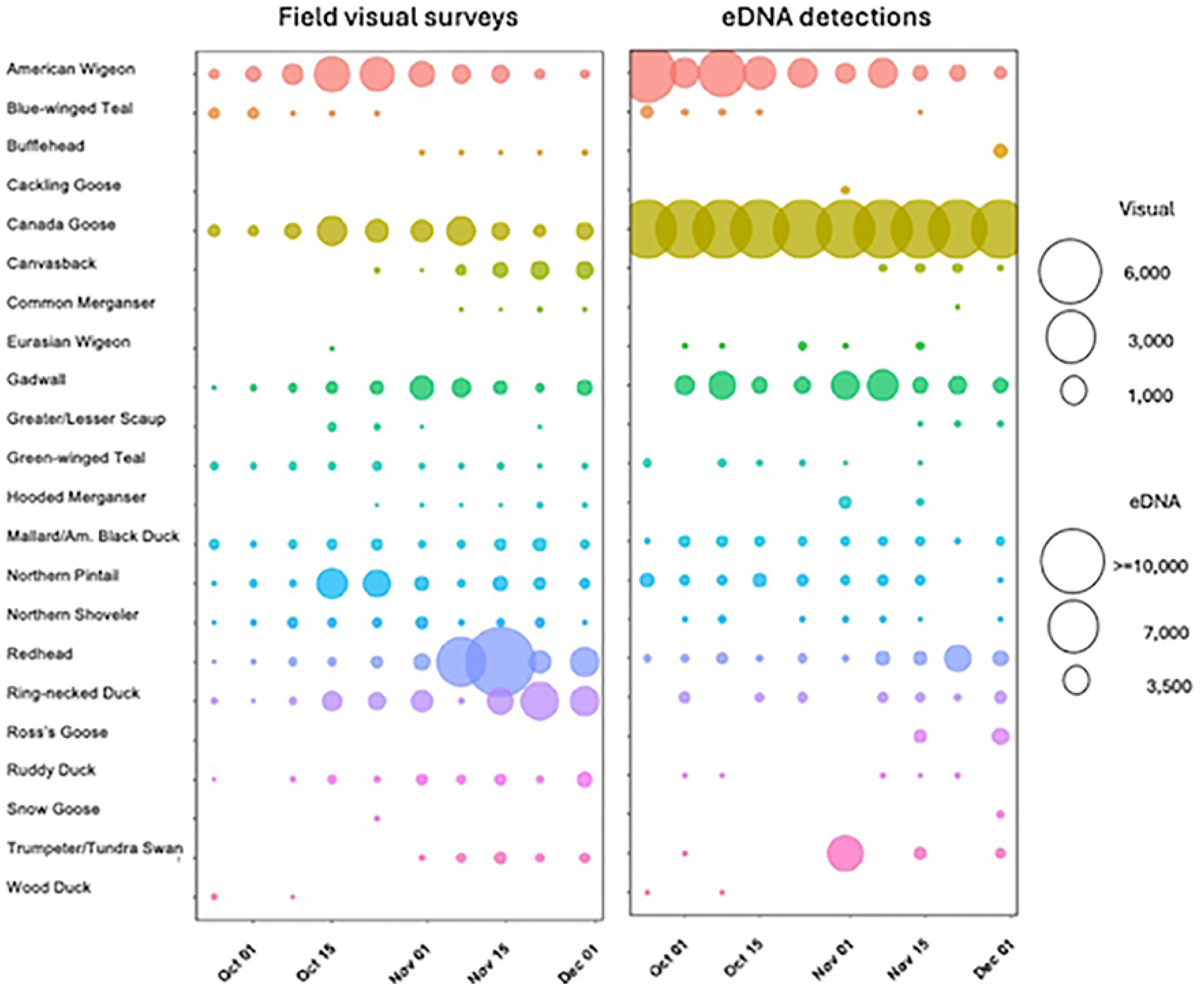
Temporal variation of visual surveys and eDNA standardized detections during waterfowl fall migration at Montezuma NWR.

In contrast, the standardized eDNA read counts showed significant correlations with relative abundance (Fig 3). A correlational network analysis of significant positive log-log relationships between ground visual surveys and standardized eDNA reads revealed a highly dynamic and variable system. Importantly, eDNA data captured relative waterfowl abundance not only on the day of sampling, but also correlated well with visual surveys conducted up to five days prior (Fig 3). The exception was October 31st, when the relative abundance of species in eDNA did not show any significant correlation with the relative abundance based on individual ground visual surveys (Fig 3).

**Fig 3.**
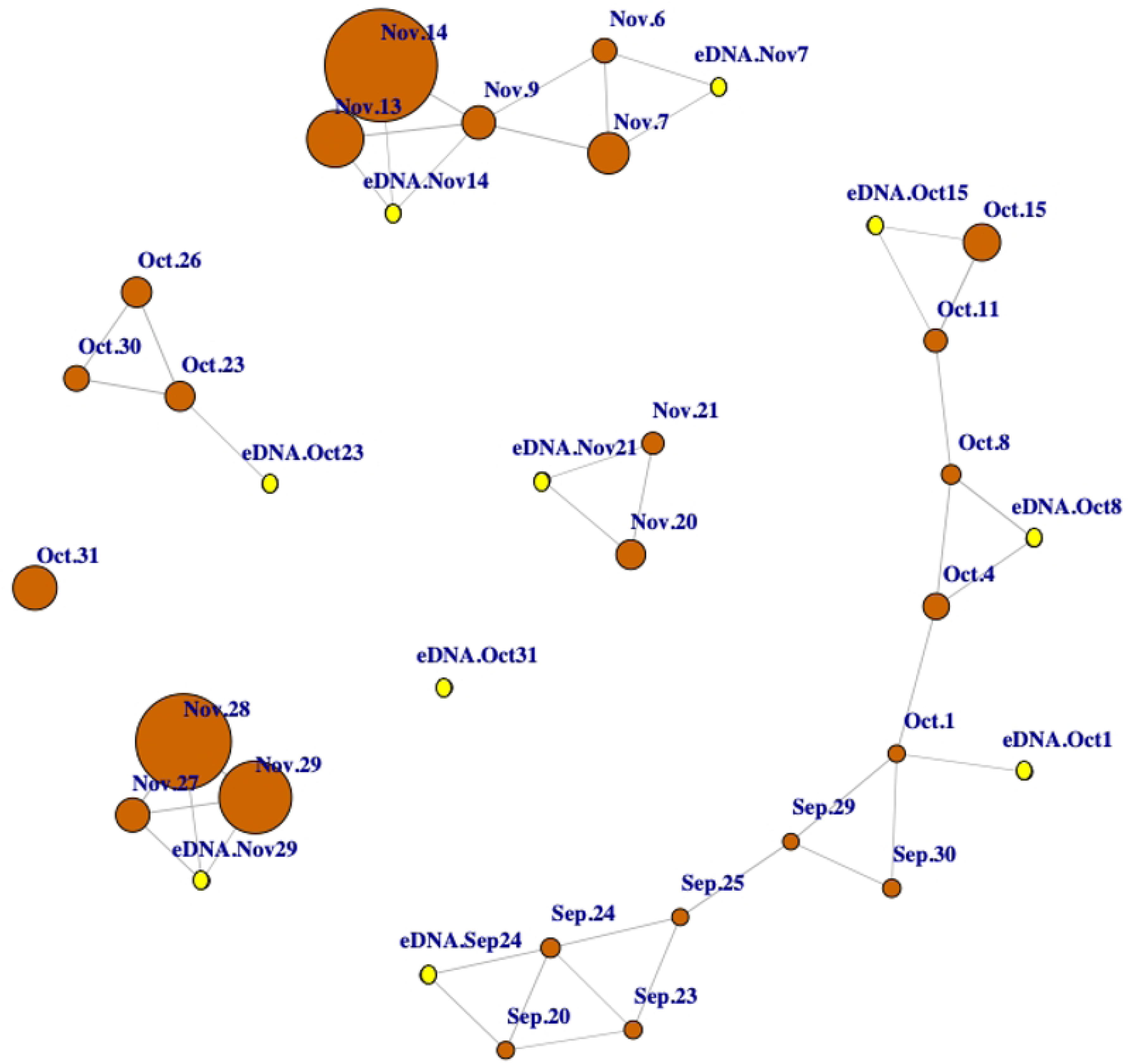
Correlation-based network of the standardized eDNA reads and the bird visual surveys. The nodes represent the dates during the waterfowl fall migration and the edges between them represent the significant, positive, and log-log correlation coefficients (≥ 0.6) between ground visual surveys (orange nodes) or between visual surveys and the standardized number of DNA reads (pink nodes) obtain by combining the Anserini/Cygnini and Anatini/Aythyini and datasets.

At any given sampling date, the relative abundance of eDNA reads showed a positive correlation (average τ ≥ 0.62) with the relative abundance of species present in the area. On 5 out of the 6 sampling dates for which at least three independent ground visual surveys (eBird) were available, the strength of the eDNA-visual correlation exceeded that of the correlation between ground visual surveys (Fig 4).

**Fig 4.**
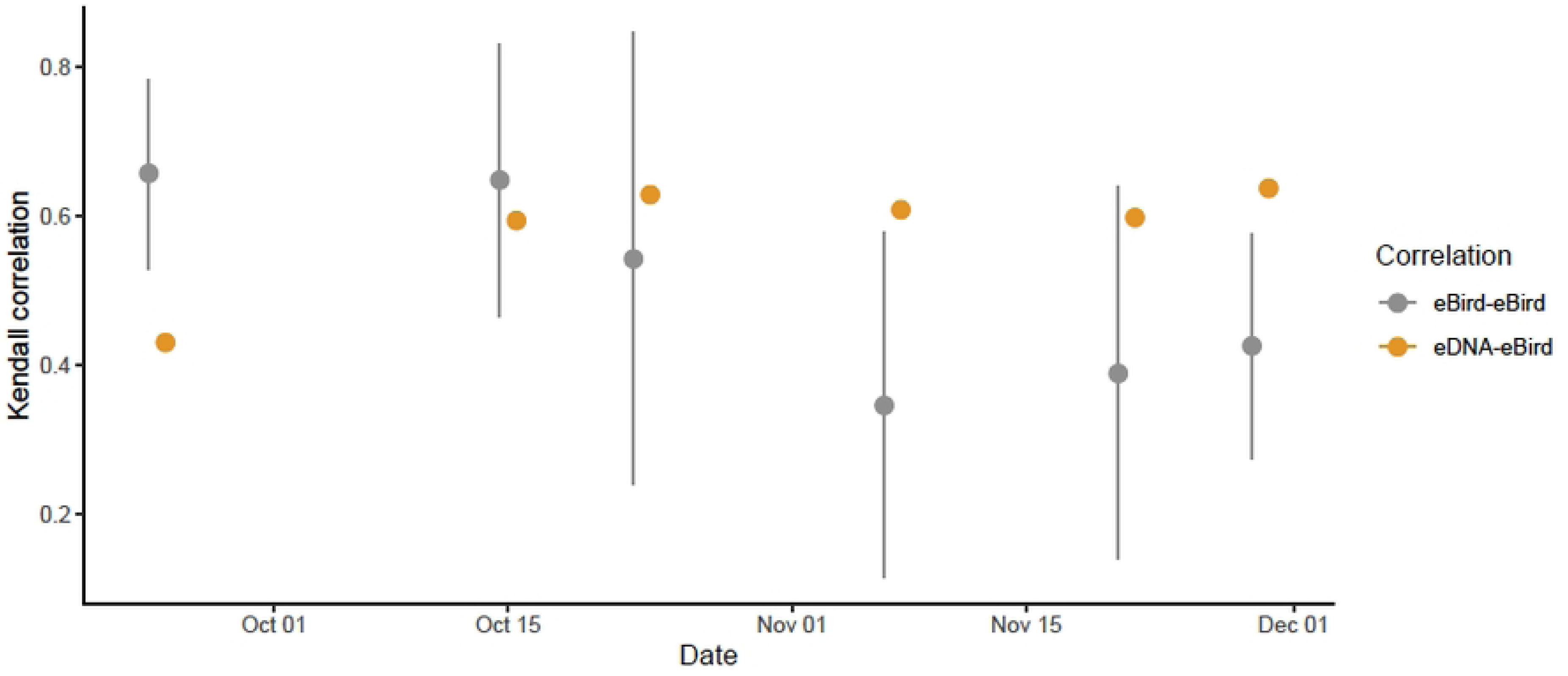
**Mean and 95% confidence intervals of Kendall correlation coefficients between eBird checklists from different observers (in gray) reported on the eDNA sampling date and between eDNA detections and eBird checklists made on the eDNA sampling date (in orange).** We did not include the dates Oct1, Oct8, Oct31 and Nov 14 in this analysis because there were fewer than three checklists available for correlation.

## Discussion

Our study adds to the growing body of research demonstrating the usefulness of eDNA approaches for monitoring bird species. While most studies have focused on the detection of specific target species—such as the Gouldian Finch [20], Black Rail [53], Ridgway’s Rail [33], and various wading birds [24, 63]—our study employs a metabarcoding approach. This method aligns with recent research exploring bird diversity in bush habitats [54], rice field landscapes [40], and waterbird communities including geese and swans [37, 64]. By employing a suite of primers specifically designed to target Anatidae tribes with reduced cross-amplification to non-target vertebrates relative to existing universal avian primers, we successfully detected both common and rare waterfowl species during the fall migration in an important bird conservation area. By collecting eDNA samples at the outflow of the refuge pool, we aimed to obtain a representative assessment of the species present in the area while minimizing disturbance. Our results indicate that this approach was effective: the environmental samples contained DNA from all waterfowl species observed during visual surveys, as well as Ross’s goose—an uncommon and difficult-to-identify species that often blends into mixed flocks with the more common snow goose.

While we successfully detected the presence of waterfowl species using eDNA, estimating their abundance remained challenging, a result that aligns with [64] using universal bird primers. We had anticipated that the use of waterfowl-specific primers would reduce PCR bias and yield more accurate estimates of individual abundance; however, this was not the case. This outcome underscores the complex relationship between eDNA concentration and organism biomass or abundance, which is influenced by species-specific DNA shedding rates, metabolic activity, and behavior [62, 79, 80]. Moreover, environmental conditions such as temperature, pH, microbial activity, and UV exposure affect DNA degradation, leading to variable eDNA persistence and detection probabilities across space and time [70, 82]. Additionally, outflow sampling is not necessarily integrating eDNA transport within the hydrodynamically complex network of interconnected pools of the refuge. All these factors may have contributed to the temporal mismatches we observed between organism and eDNA abundance.

However, the proportion of sequence reads attributed to different waterfowl species reliably reflected their relative abundance. This approach enabled us to detect shifts in the waterfowl community composition during the fall migration period, providing valuable ecological insights without the need for precise population counts. By observing the variation in relative abundance of waterfowl species with eDNA, we learned that the waterfowl community at Montezuma NWR is a highly dynamic system during the fall migration, with changes in species composition. Using eDNA, we were able to track variation in species presence over time quite well for some species, e.g., canvasback, wood duck, and blue-winged teal. The utility of relative abundance metrics is increasingly recognized, particularly for tracking phenological and distributional changes in migratory birds [25, 29, 30, 44]. Long-term avian monitoring programs have shown that shifts in relative abundance are often early indicators of habitat degradation, climate-driven range shifts, and biodiversity loss [12, 36, 61]. For example, North America bird abundance has declined since 1970, with grassland and migratory species showing the steepest losses—a trend that was initially detected through changes in relative species occurrence and reporting frequency [61]. Although waterfowl populations have shown relative resilience and even growth, making them one of the few avian guilds with increasing or stable trends [74] shifts in the relative abundance of dabbling ducks versus diving ducks in wetland systems can indicate ecological changes such as alterations in vegetation structure or water depth, which are key factors for guiding adaptive management strategies [8].

In our study, eDNA detections, similarly to our visual surveys, also demonstrated shifts in the relative abundance for some dabbling ducks, more abundant between early and mid-fall migration (for example, Blue-winged Teal and Wood Duck) and diving ducks, more abundant during late fall migration (for example, Canvasback and Redhead). Over longer timescales, consistent tracking of the relative abundance of waterfowl using the approach described in this paper can also reveal the cumulative impacts of conservation interventions, including wetland restoration and adaptive water management aimed at enhancing conditions for target species [8, 46].

The relative abundance of eDNA reads showed a consistently positive correlation with the observed species composition, and the correlation coefficients between eDNA and ground visual surveys were stronger than the correlation among the ground-based surveys themselves. These results suggest that eDNA-targeted metabarcoding not only captures ecologically meaningful signals of relative abundance but may also reduce the variability and subjectivity inherent in traditional methods. While eDNA-targeted metabarcoding is not yet a replacement for traditional monitoring methods, it serves as a powerful complementary tool for informing adaptive management in dynamic wetland ecosystems facing increasing environmental pressures.

## Acknowledgments

We thank Matthew Medler (Cornell Lab of Ornithology, Cornell University) for support to L. Andrade during the visual bird surveys and field sampling and for reviewing the manuscript. We also thank Linda Ziemba (Wildlife Biologist, Montezuma NWR) for all her support in the field sampling at Montezuma NWR, and Anne Rhoads for all her support during the field work at the Huyck Preserve. We thank Irby J. Lovette, Bronwyn Butcher, and the Lovette Lab for all their support to L. Andrade with the pilot molecular analyses. We also would like to thank Kara Andres, Paul Simonin, and Timothy Lambert for feedback on the field sampling and molecular analyses.

## Supporting information

**S1 Fig. eDNA sampling sites at the Cornell Lab of Ornithology, Ithaca, NY (A), sampling site SWP1, and the Huyck Preserve, Rensselaerville, NY (B). Lake Lincoln Pond (P1 and P2) and Lake Myosotis (L1 – L3).Ten Mile Creek (C1), which links the two waterbodies.**

**S2 Fig. Workflow of the pilot experiments to conduct the initial assessment of both general and targeted bird metabarcoding primers (the Cornell Lab of Ornithology, Phase I, and the Huyck Preserve, Phase II).**

**S3 Fig. Percentage of sequence reads per organism group for each genetic marker tested to amplify bird DNA in environmental samples collected at Sapsucker Woods Pond and Huyck Preserve.**

**S4 Fig. Phylogenetic tree of waterfowl species selected in this study to design waterfowl ND2 primers.** We used the Neighbor-Joining tree-built method and Tamura-Nei genetic distance model in Geneious Prime 2020.2.1. Colors represent different Tribes: red (Anserini and Cygnini), green (Anatini, Cairinini, and Oxyura), and orange (Aythyini and Mergini).

**S5 Fig. Graphical summary of the taxonomic families amplified by waterfowl ND2 primers.** (A) Anatini and Cairinini ND2 Primers, (B) Aythyini and Mergini ND2 Primers, and (C) Anserini and Cygnini ND2 Primers.

**S1 Table. Mitochondrial genes and multiple primers tested in the pilot experiments to amplify bird DNA from eDNA water samples.** Different colors in the table represent the primer pair. Mibird-U-F1, F2, F3, and F4 were used in pair with Mibird-U-R.

**S2 Table. Insert length and amplicon size of each genetic marker tested in this study. S1 File. Supporting information**

